# Modular transcriptional programs separately define axon and dendrite connectivity

**DOI:** 10.1101/724633

**Authors:** Yerbol Z. Kurmangaliyev, Juyoun Yoo, Samuel A. LoCascio, S. Lawrence Zipursky

## Abstract

Patterns of synaptic connectivity are remarkably precise and complex. Single-cell RNA sequencing has revealed a vast transcriptional diversity of neurons. Nevertheless, a clear logic underlying the transcriptional control of neuronal connectivity has yet to emerge. Here, we focused on Drosophila T4/T5 neurons, a class of closely related neuronal subtypes with different wiring patterns. Eight subtypes of T4/T5 neurons are defined by combinations of two patterns of dendritic inputs and four patterns of axonal outputs. Single-cell profiling during development revealed distinct transcriptional programs defining each dendrite and axon wiring pattern. These programs were defined by the expression of a few transcription factors and different combinations of cell surface proteins. Gain and loss of function studies provide evidence for independent control of different wiring features. We propose that modular transcriptional programs for distinct wiring features are assembled in different combinations to generate diverse patterns of neuronal connectivity.

## Introduction

Brain function relies on precise patterns of synaptic connections between neurons. At the cellular level, this entails each neuron adopting a specific wiring pattern, the combination of specific synaptic inputs and outputs. In invertebrates, stereotypical wiring patterns are genetically encoded in the programs regulating the development of neurons. Much of the specificity of inputs and outputs of neurons in the mammalian CNS is also genetically determined (Sanes and Zipursky, 2010).

Vast numbers of neurites from a diversity of neurons are intermingled within the developing central nervous system, and they form highly specific synaptic connections with a discrete subset of the neurons they contact. Studies in both vertebrates and invertebrates have led to the identification of cell surface proteins (CSPs) that mediate selective association between neurites (Tessier-Lavigne and Goodman 1996; de Wit and Ghosh, 2016; Zinn and Ozkan, 2017). Gain and loss of function genetic studies have shown that combinations of different CSP families regulate this specificity (Zarin et al. 2014). Indeed, neuronal subtypes express highly diverse repertoires of CSPs during circuit assembly (Tan et al., 2015; Li et al., 2017; Sarin et al., 2018). Conserved regulatory strategies involving combinations of transcription factors (TFs) establish unique neuronal identities (Allan and Thor, 2015; Enriquez et al., 2015; Hobert, 2016). However, the programs regulating expression of CSPs for specific neuronal wiring features are still poorly understood.

Single-cell RNA sequencing (RNA-Seq) provides an unsupervised approach to uncover the genetic programs underlying specific wiring features by exploring subtype-specific transcriptomes during development. As neuronal subtypes exhibit differences in characteristics other than wiring patterns, the relationship between genes and wiring specificity may be obscured by genes contributing to other aspects of neuronal diversity. Therefore, sets of closely related neurons with highly specific differences in wiring patterns are ideally suited to uncover the genetic programs specific to wiring. Here, we explore the genetic logic underlying synaptic specificity in one such set of neurons: T4/T5 neurons of the *Drosophila* visual motion detection pathway. We envision that our findings in this system will provide insights into the genetic logic of wiring specificity more broadly in both vertebrate and invertebrate systems.

T4/T5 neurons share a common developmental origin, physiological function, and morphology, but differ in their precise wiring patterns and preferred stimulus (Fischbach and Dittrich, 1989; Maisak et al., 2013; Apitz and Salecker, 2018; Pinto-Teixeira et al., 2018; Shinomiya et al., 2019). T4/T5s can be classified into two quartets of subtypes based on dendritic inputs: the four T4 subtypes share a common set of dendritic inputs in the medulla, and the four T5 neurons share a different set of dendritic inputs in the lobula (Figure 1a-c). T4 neurons respond to ON stimuli (i.e. bright edges moving against a dark background) and T5 to OFF stimuli (i.e. dark edges moving across a bright background). T4/T5 neurons can also be classified into four pairs of subtypes (a-d) based on the location of their axon terminals in layers a-d of the lobula plate (LoP). Each pair responds selectively to visual motion in one of four cardinal directions: posterior, anterior, upwards, and downwards motion, respectively (Figure 1a-c). We hypothesized that comparisons between the gene expression patterns of developing T4/T5 subtypes would provide insight into the genetic programs regulating discrete wiring features.

**Figure 1.**
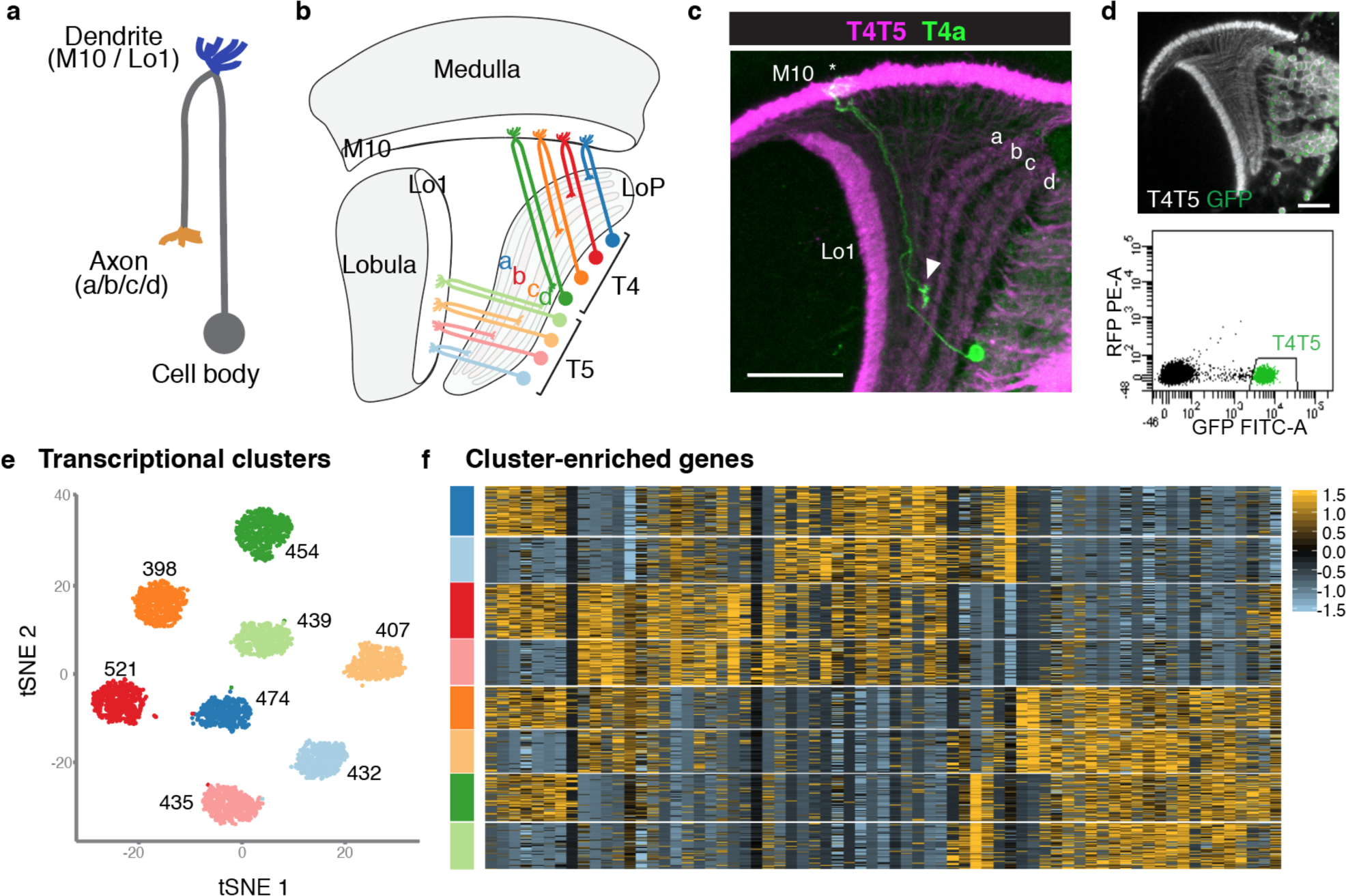
Single-cell sequencing reveals eight transcriptionally distinct populations of T4/T5 neurons. (a) Common morphology of a T4/T5 neuron, with axon and dendrite wiring pattern variations in parentheses. (b) Arrangement of the eight T4/T5 subtypes in the optic lobe. Each subtype is defined by a combination of one dendrite (M10 or Lo1) and one axon (LoP a, b, c, or d) wiring pattern. (c) A single T4a neuron (green) with dendrites in M10 (asterisk) and axon terminal in LoP layer a (arrowhead). All T4/T5 neurons labeled in magenta. Scale bar, 20μm. (d-f) Single-cell sequencing of T4/T5 neurons at 48h APF. Unsupervised analysis revealed eight distinct transcriptional clusters. (d) T4/T5 neurons were labeled with nuclear GFP, purified by FACS and used for single-cell RNA-Seq. (e) t-distributed stochastic neighbor embedding (tSNE) plot of 3557 single-cell transcriptomes. Clusters are color-coded according to subtype identity based on following results. Cell numbers are displayed for each cluster. (f) Heatmap of expression patterns of cluster-enriched genes (“one versus all”, see Methods). Cells (rows) grouped by cluster identities as in (e). Genes (columns) are ordered by similarity of their expression patterns. Scaled expression levels are indicated, as in scale.

Here, we report that independent transcriptional programs define the dendritic inputs and axonal outputs of T4/T5 neurons. We present gain and loss of function studies indicating that these programs control their corresponding morphological features. Our findings suggest that the modular assembly of separate dendritic and axonal transcriptional programs contributes to the diversity of wiring patterns in complex nervous systems.

## Results

### Single-cell RNA-Seq reveals eight transcriptionally distinct populations of T4/T5 cells

As a step towards uncovering genetic programs that control neuronal wiring patterns, we performed single-cell RNA-Seq on developing T4/T5 neurons. Sequencing was performed at 48h after puparium formation (APF). This developmental time point precedes a period of widespread synaptogenesis in the visual system, and coincides with the appearance of four discrete synaptic layers (a, b, c, d) in the LoP neuropil. Neurons were purified from dissected optic lobes by FACS using a transgenic line with nuclear GFP selectively expressed in all T4/T5 neurons (Figure 1d). RNA-Seq libraries were generated using 10X Chromium technology (Zheng et al., 2017) and sequenced to a mean depth of 92,000 reads per cell. In total, we profiled 3894 cells with a median of 1633 genes and 4389 transcripts captured per cell. After quality control and removal of outlier cells, our final dataset consisted of 3557 cells with 1000-2000 genes per cell.

We applied independent component analysis (ICA) followed by a graph-based clustering method to separate transcriptionally distinct cell populations (Butler et al., 2018; Saunders et al., 2018). Unsupervised analysis revealed eight clusters of approximately equal numbers of cells (Figures 1e and Supplementary Figure 1), suggesting that each cluster corresponded to a single T4/T5 subtype.

To identify genes preferentially expressed in T4/T5 subtypes, we performed differential gene expression analysis between each of the eight individual clusters and all other cells in the dataset (i.e. “one versus all,” see Methods). This revealed 69 genes which were strongly expressed in some clusters and not in others. Cluster-enriched genes, however, were not specific to single clusters. By contrast, for instance, each of the five subtypes of lamina neurons is defined by at least one subtype-type specific transcription factor (Tan et al. 2015). Thus, while T4/T5 subtypes separated into eight transcriptionally distinct clusters, they were not defined by unique molecular markers (Figure 1f).

### Eight T4/T5 transcriptional clusters are separated by three primary axes of transcriptional diversity

The absence of unique markers for individual subtypes suggested that they were instead defined by unique combinations of genes. ICA has been shown to capture groups of genes corresponding to discrete biological phenomena (Saunders et al., 2018). Intriguingly, three independent components (ICs) each split the eight T4/T5 clusters into two groups of four, each in a different way (Figure 2a). Together, these three ICs were sufficient to define all eight clusters (Figure 2b).

**Figure 2.**
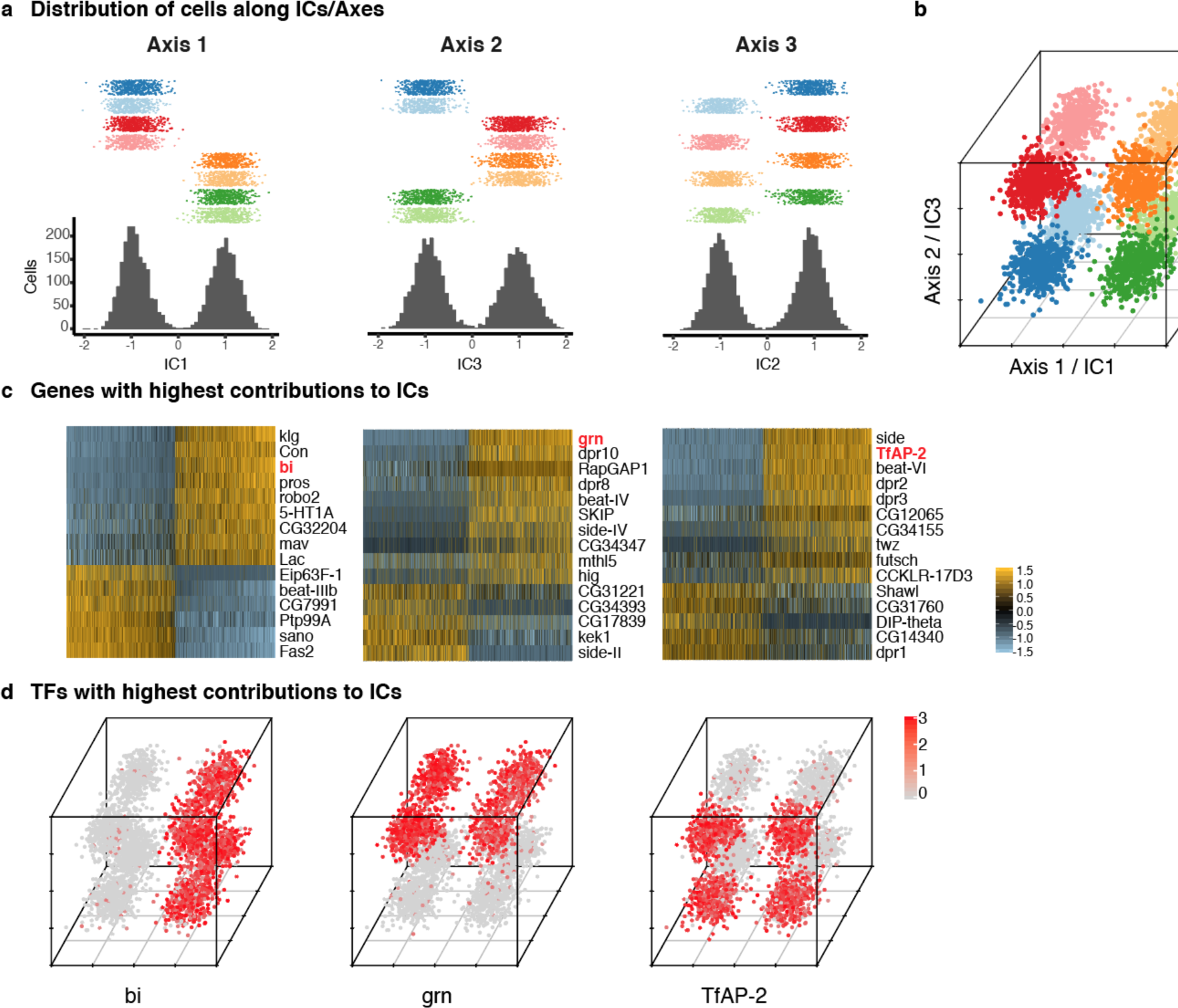
Three primary axes of transcriptional diversity define eight T4/T5 populations. (a) Three independent components (ICs, henceforth Axis 1, 2, 3) separate cells into approximate halves. Histograms (bottom) and 1-D scatterplots (top) show the distributions of cells along each axis. Cells are grouped into rows based on cluster identities. ICs/Axes are ordered according to following results. Clusters are color-coded as in Figure 1e. See also Supplementary Figure 1. (b) 3-D scatterplot of the distributions of cells along the three ICs/Axes. (c) Heatmaps of expression patterns of the top 15 genes with highest contribution (loading) to each IC/Axis. Cells (columns) are ordered according to a score for each IC/Axis. Genes (rows) are ordered according to the contribution to each IC/Axis. Scaled expression levels are indicated, as in scale. Axes ordered as in (a). (d) 3-D scatterplots with expression patterns of transcription factors (TFs) with highest contribution to each IC/Axis. Normalized expression levels are indicated by color, as in scale. Axes are arranged as in (b).

Each of the three ICs defined an axis of transcriptional diversity (hereafter referred to as Axis 1, 2, 3) driven by a group of genes differentially expressed along each axis. Many of these genes were expressed in a binary (ON/OFF) pattern in one of the two groups of clusters separated along each axis (Figure 2c). A small set of TFs were among the genes with the highest contributions to each axis and illustrate this pattern. Binary expression of *bifid* (*bi*) defined the two groups of clusters separated by Axis 1, *grain* (*grn*) defined the clusters separated by Axis 2, and *TfAP-2* defined the clusters separated by Axis 3 (Figure 2d). Thus, while no individual cluster is uniquely defined by the expression of a single gene, each cluster expresses a unique combination of genes. In this way, three axes of diversity with orthogonal ON/OFF expression patterns of TFs define the eight T4/T5 clusters.

### Primary axes of transcriptional diversity correspond to axon and dendrite wiring patterns

We next sought to map transcriptional clusters to T4/T5 subtypes, and to determine the biological significance of the observed axes of transcriptional diversity. We inspected *in vivo* expression patterns of genes associated with the three primary axes of diversity using transgenic reporters inserted into the endogenous loci (Venken et al., 2011).

Axis 1 separated clusters into two groups of four that were defined by mutually exclusive binary expression of two genes, *Fasciclin 2* (*Fas2*) and *klingon* (*klg*), respectively (Figure 3a), each encoding immunoglobulin (Ig) superfamily proteins. *Fas2* was expressed in LoP layers a/b, whereas *klg* was expressed in LoP layers c/d (Figure 3b). Clusters expressing *Fas2* and *klg* also expressed previously described markers for T4/T5 subtypes a/b (*dachshund* (*dac*)) and c/d (*bi*, *Connectin* (*Con*)) (Apitz and Salecker, 2018) (Supplementary Figure 2). Thus, Axis 1 separated LoP layer a/b and c/d subtypes, defining specificity of axonal outputs between two broad domains of the LoP (Figure 3c). This corresponds to separation of horizontal (posterior/anterior) and vertical (upwards/downwards) motion detection circuits.

**Figure 3.**
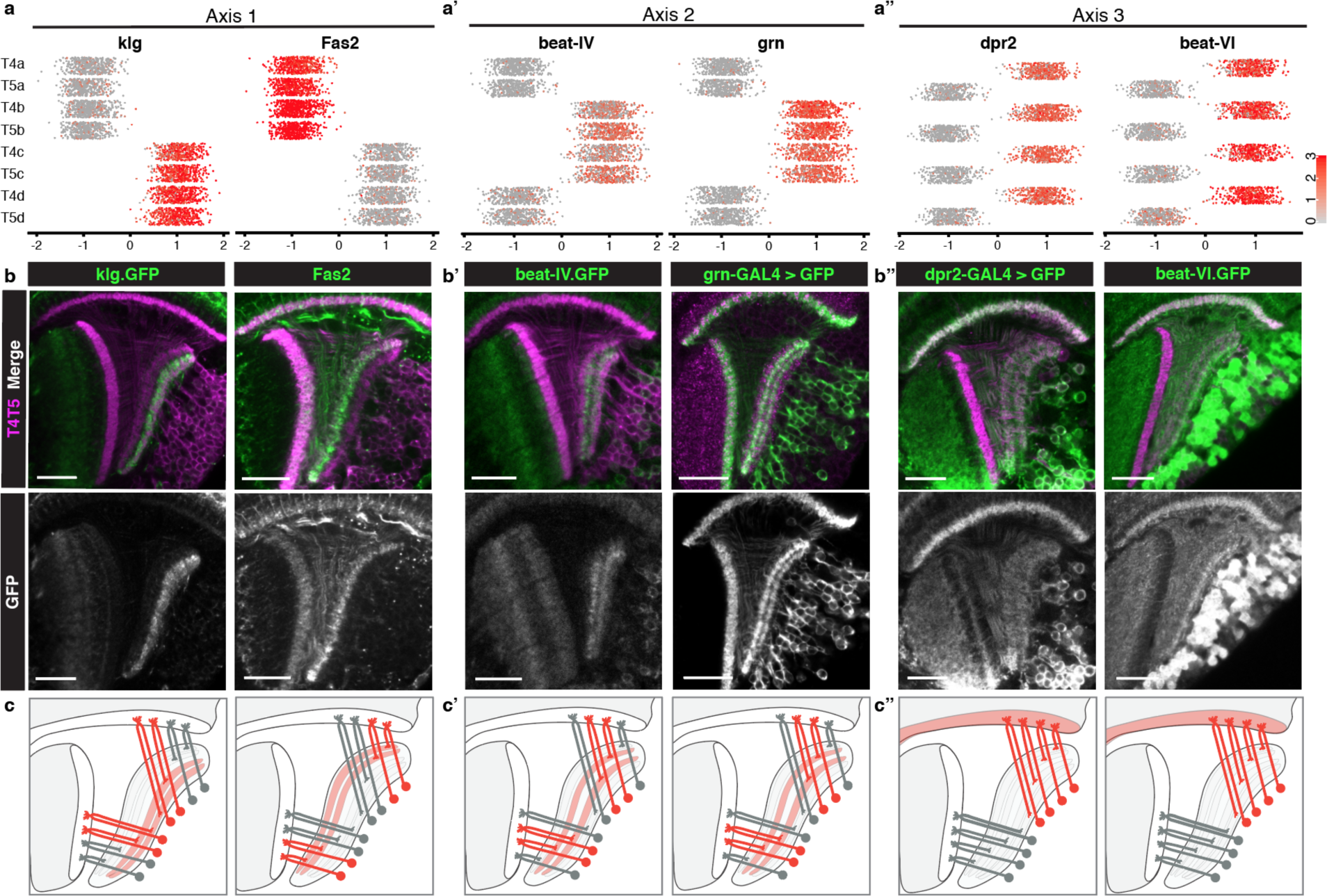
Primary axes of transcriptional diversity define groups of T4/T5 subtypes with shared wiring patterns. (a-a”) 1-D scatterplots show distribution of cells along Axis 1, 2, and 3 for each cluster. Normalized expression levels are indicated by color, as in scale. (b-b”) *In vivo* expression of marker genes for each axis at 48h APF. *klg* labels LoP layers c/d, *Fas2* labels LoP layers a/b, *beat-IV* and *grn* label LoP layers b/c, *dpr2* and *beat-VI* label M10 but not Lo1. Scale bars, 20μm. Sets of positive clusters in (a) are matched to specific sets of T4/T5 subtypes based on *in vivo* expression patterns in (b). Individual cluster identities are deduced based on combination of expression patterns. For example, T4a is *Fas2+* (a/b), *beat-IV-*(not b/c), *dpr2+* (M10). (c-c”) Schematic of wiring patterns of T4/T5 subtypes corresponding to the expression patterns of marker genes (red).

Axis 2 separated clusters into two groups of four defined by binary expression of *beat-IV* (an Ig superfamily protein) and the TF *grn*. Both genes were expressed in LoP layers b/c, but not a/d. Thus, Axis 2 further separated subtypes into inner (b and c) and outer (a and d) LoP layer subtypes in a symmetrical fashion, defining specificity of axonal outputs between adjacent layers within the two broad domains of the LoP (Figure 3a’-3c’). This corresponds to further separation of each of the motion detection circuits into two subcircuits detecting motion in two opposing directions (i.e. horizontal into posterior and anterior, and vertical into upwards and downwards motion).

Axis 3 separated clusters into two groups of four defined by binary expression of *dpr2* and *beat-VI*, each encoding an Ig superfamily protein. *In vivo*, both genes were expressed in all LoP layers, and M10 but not Lo1. Thus, Axis 3 separated all T4 from all T5 subtypes, defining specificity of dendritic inputs (Figure 3a’’-3c’’). This corresponds to separation between inputs from two parallel circuits upstream of T4/T5 neurons (ON and OFF motion detection pathways).

Taken together, three primary axes of diversity defined distinct wiring features of T4/T5 subtypes and in combination defined wiring patterns of each T4/T5 subtype (Figure 3a-3a’’). A combination of Axis 1 and Axis 2 defined four types of axonal outputs (a, b, c, d), and Axis 3 defined two types of dendritic inputs (T4 and T5).

### Transcriptional program of a single T4/T5 subtype

In addition to the transcriptional differences between groups of T4/T5 subtypes described above, further variation might exist at the individual subtype level. To examine this question, we focused on a single subtype (T4a) and performed comprehensive pairwise comparisons with each of the other subtypes (i.e. “one versus one”).

First, we compared T4a and each subtype that differed by a single wiring feature: either axonal outputs (T4b, T4c, T4d), or dendritic inputs (T5a) (Figure 4, upper dot plots). Comparison of T4a to T4c or T4d, which have axonal outputs in non-adjacent LoP layers, yielded the largest number of DEGs (58 and 55, respectively). T4a and T4b have axonal outputs in adjacent LoP layers, and were separated by an intermediate number of DEGs (16). Finally, only a small number of DEGs (9) separated T4a and T5a, which share axonal output but receive different dendritic inputs.

**Figure 4.**
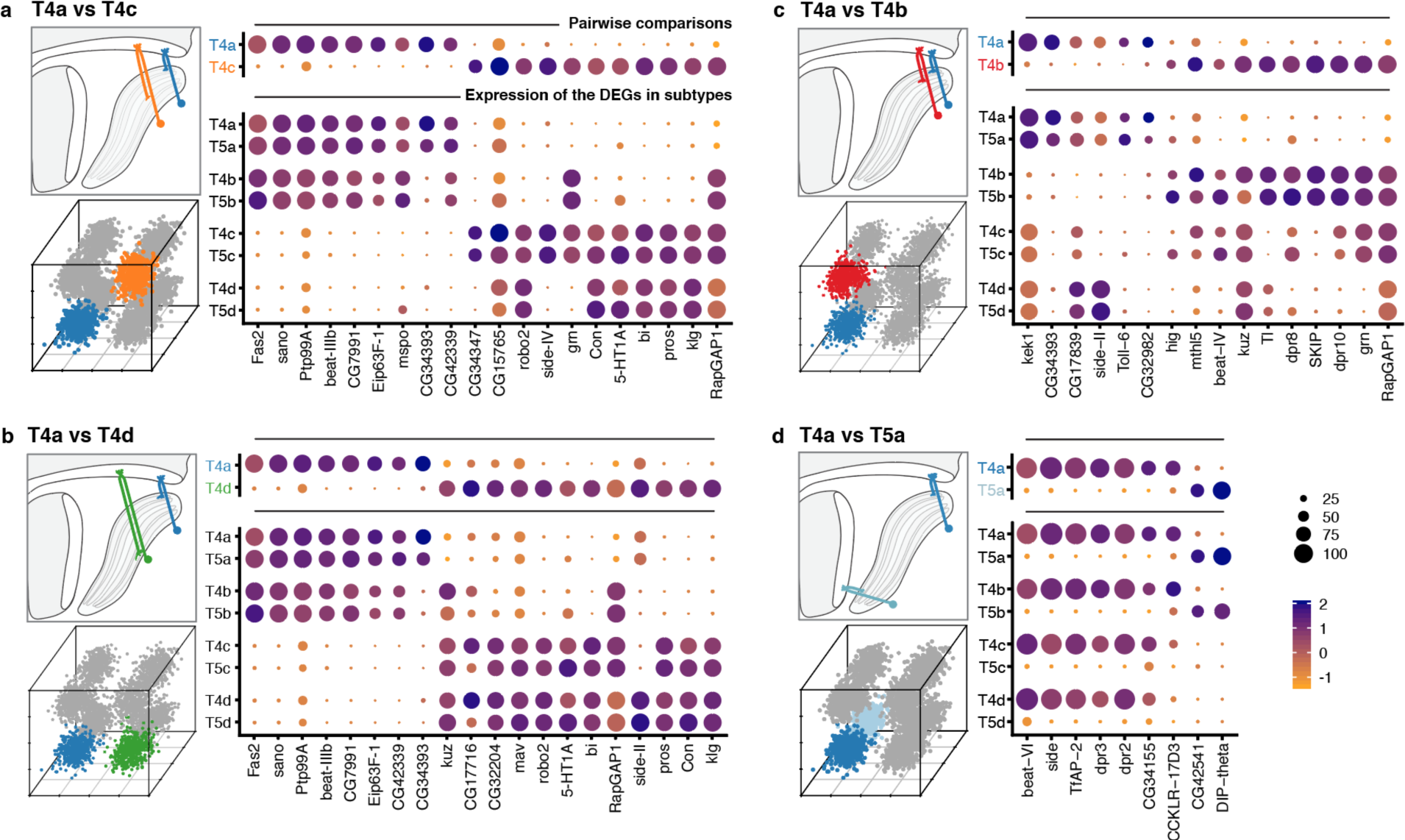
Transcriptional program of a single T4/T5 subtype. Pairwise comparisons between T4a and other subtypes (“one versus one”, see Methods) that differ by either axonal outputs (a-c), or dendritic inputs (d). For each comparison, insets indicate morphologies (upper left) and cluster distributions along axes of transcriptional diversity (lower left). Expression patterns of differentially expressed genes (DEGs) for each pairwise comparison are shown in upper right. Dot size indicates the percentage of cells in which the DEG was detected, color represents average scaled expression, as in scale. Genes are ordered by fold-change values. Top 20 DEGs are shown for (a) and (b). Expression patterns of DEGs among all eight subtypes are shown in lower right.

Expression patterns of the DEGs from pairwise comparisons across all T4/T5 subtypes revealed a general pattern (Figure 4, lower dot plots): virtually all DEGs were co-regulated across all subtypes according to either specificity of axonal outputs or dendritic inputs. In other words, distinct sets of DEGs were expressed in each pair of subtypes with shared axonal outputs, but different dendritic inputs (e.g. in T4a and T5a, Figure 4a-c). Similarly, a distinct set of DEGs was expressed in groups of subtypes with shared dendritic inputs, but different axonal outputs (i.e. all T4 subtypes, Figure 4d).

The co-regulation of DEGs according to wiring patterns was not limited to comparisons between subtypes that differed by a single wiring feature. DEGs between T4a and subtypes that differed by both axon and dendrite wiring patterns (e.g. T5c), were also expressed in groups of subtypes sharing either axonal outputs or dendritic inputs (Supplementary Figure 3).

In addition to three primary axes of transcriptional diversity, this analysis shows that a number of DEGs exhibited more distinct LoP layer-specific patterns. For example, many DEGs were specifically expressed or suppressed in T4/T5 subtypes from a single LoP layer (Figure 4c).

Many of the DEGs have been implicated in neuronal wiring specificity (Figure 4). Approximately half of the DEGs encoded CSPs with cell adhesion domains (Ig/LRR), including multiple members of the dpr/DIP and beat/side families of interacting proteins (Zinn and Özkan, 2017); specific members of these families have been shown to regulate axon guidance and synaptic specificity in the developing fly nervous system.

Taken together, these results reveal that the transcriptional organization of T4/T5 neurons mirrored their wiring patterns. Discrete groups of co-regulated genes reiteratively defined either shared axon or shared dendrite wiring patterns among different subtypes. These groups of genes were assembled in different combinations to uniquely define the eight T4/T5 subtypes.

### Stable and dynamic features of T4/T5 transcriptional programs during development

To evaluate how gene expression in T4/T5 subtypes changes during development, we profiled T4/T5s at an earlier time point, 24h APF. Similar to our dataset at 48h APF, we identified eight distinct populations separated by three equivalent axes of transcriptional diversity. Many of the same genes were associated with these axes at both time points, allowing us to match subtypes between 24h and 48h APF (Supplementary Figure 4).

Comparison of 24h and 48h datasets revealed stable and dynamic features of T4/T5 transcriptional programs. TFs defining the primary axes of diversity (*bi, grn, TfAP-2*) were expressed in the same sets of subtypes at both time points, suggesting they may contribute to stable subtype identities during development (Figure 5a). Some CSPs also exhibited stable expression, marking subtypes with shared axon or dendrite wiring patterns at both time points. Other CSPs were dynamically regulated and were specific to subtypes only at a particular stage of development (Figure 5b-d).

**Figure 5.**
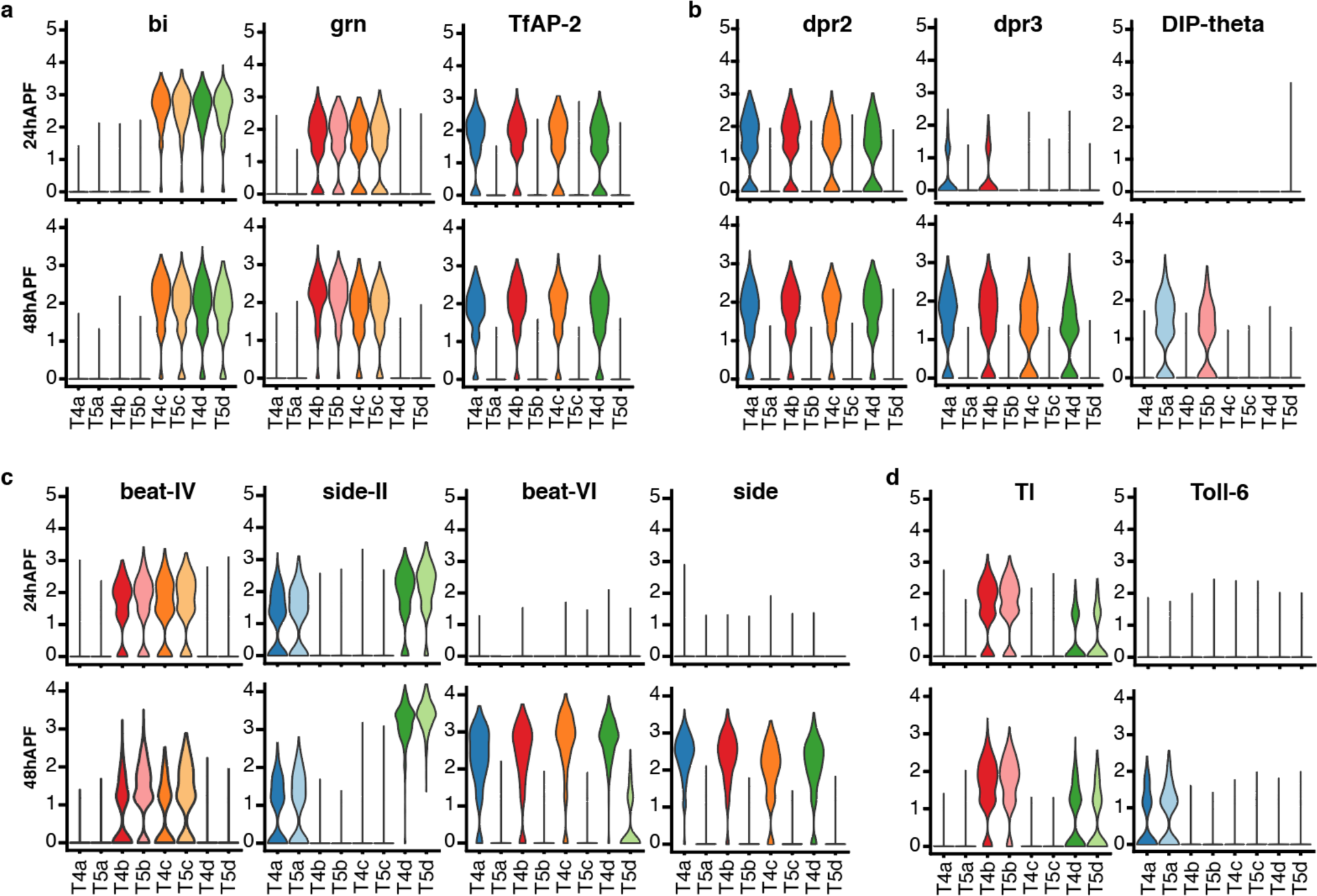
Dynamics of T4/T5 transcriptional programs during development. Distributions of normalized expression levels of TFs and selected families of CSPs at 24h and 48h APF. Distributions for each subtype are color-coded as in Figure 1. See also Supplementary Figure 4.

Interestingly, dynamic changes in gene expression were also coordinated among subtypes with shared wiring features (Figure 5b-d). For example, *dpr3* and a few other CSPs were synchronously upregulated in all T4 subtypes from 24h to 48h APF. Similarly, *Toll-6* was synchronously upregulated in both LoP layer “a” subtypes (T4a/T5a). These data indicate that similar transcriptional programs unfold in parallel among T4/T5 subtypes with shared wiring features.

### Axon-specific transcriptional programs of T4/T5 neurons control lamination of LoP layers

A remarkable resemblance between transcriptional programs and wiring patterns suggested that these programs control development of corresponding features of T4/T5 neurons. During the period covered in our study (24 - 48h APF) four LoP layers form in two discrete lamination steps (Supplementary Figure 5). The T4/T5 axon terminals first laminate into two broad domains corresponding to layers a/b and c/d. These domains then further sublaminate into two pairs of adjacent layers to form the four discrete LoP layers, a, b, c, and d. The two primary axes of transcriptional diversity mirrored these two stages of LoP layer formation, suggesting a regulatory code for axon wiring (Figure 3). Mutually exclusive expression of TFs *dac* and *bi/pros* separated a/b (*dac*+) from c/d (*bi*+/*pros*+) subtypes (i.e. Axis 1). These subtypes were further separated by expression of the TF *grn* into inner (b and c, *grn*+) and outer (a and d, *grn-*) layer subtypes in a symmetrical fashion (i.e. Axis 2). This suggested that two levels of transcriptional regulation, acting either sequentially or in a temporally overlapping way, control development of four types of T4/T5 axonal outputs.

We sought to experimentally address this issue. Previous studies indicate that *bi* specifies c/d subtypes and formation of corresponding LoP layers. RNAi of *bi* in all T4/T5 neurons results in loss of the c/d domain of the LoP, whereas overexpression results in loss of the a/b domain. In both cases, further sublamination of remaining inner and outer LoP layer pairs still occurs (Apitz and Salecker, 2018). We performed RNAi of *grn*, which resulted in a different phenotype; whereas distinct a/b and c/d LoP domains were still separated by a pronounced gap and differential expression of *Con* (a marker for LoP layers c/d), both domains failed to sublaminate into inner and outer layers, instead forming a single layer each (Figure 6a-b). The overall morphological organization of T4/T5 neurons was otherwise unaffected. Overexpression of *grn* in all T4/T5 neurons also resulted in a specific failure of a/b and c/d LoP domains to sublaminate.

**Figure 6.**
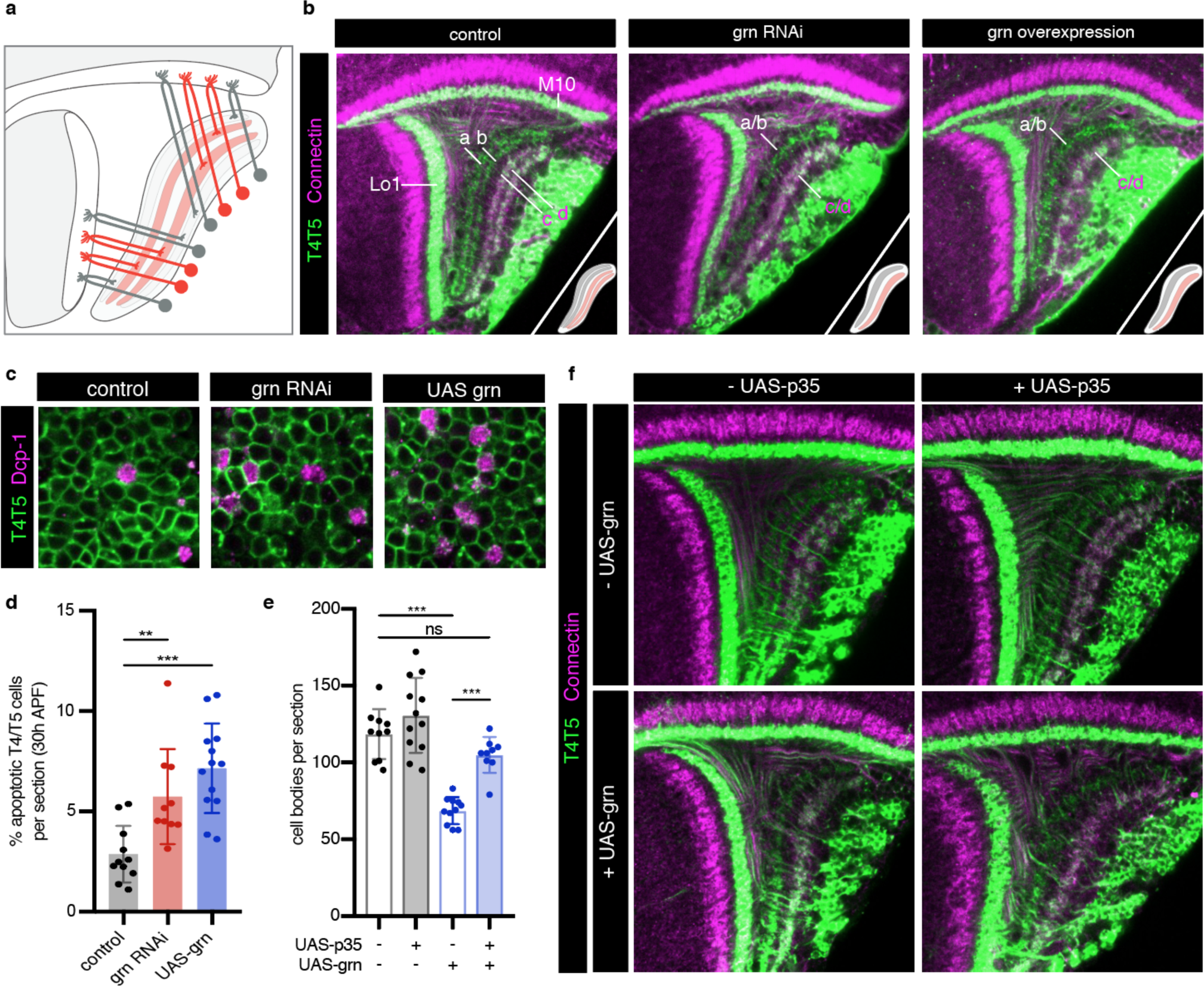
*grn* controls sublamination of T4/T5 axons into inner and outer LoP layers. (a) Schematic of *grn*+ (red) T4/T5 subtypes in wild-type optic lobe. *grn* expression defines inner LoP layer subtypes. (b) *grn* RNAi and *grn* overexpression in all T4/T5 neurons specifically disrupts sublamination of a/b (Con-) and c/d (Con+) subdomains into inner and outer layers. Insets depict LoP phenotypes. (c-d) Immunostaining for Death caspase 1 (Dcp-1) reveals increased apoptotic T4/T5 neurons under *grn* RNAi and overexpression (UAS-grn) conditions at 30h APF. See also Supplementary Figure 6. (e) Ectopic expression of p35 in T4/T5 neurons (UAS-p35) rescues apoptotic cell death associated with overexpression of *grn*. (f) *grn* overexpression specifically disrupts axon sublamination when apoptosis is blocked. Statistical significance assessed by one-way ANOVA with Tukey’s multiple comparison test (**p<0.01, ***p<0.001). Bars and whiskers represent mean and standard deviation. Dots represent values for individual optic lobes.

RNAi and overexpression of *grn* resulted in significant loss of T4/T5 neuron numbers between 24h and 48h APF (Supplementary Figure 6), associated with an increase in apoptosis (Figure 6c-d). Failure of LoP layer sublamination could result from death of specific subtypes during development. Alternatively, differential expression of *grn* might be required to direct T4/T5 axons to discrete layers. Expression of baculovirus caspase inhibitor p35 in developing T4/T5s rescued cell death associated with *grn* overexpression. Nevertheless, T4/T5 axons still failed to sublaminate into four discrete LoP layers (Figure 6e-f). Thus, *grn* is specifically required for sublamination of T4/T5 axons between pairs of adjacent layers of LoP.

We conclude that axon-specific transcriptional programs defined by binary (ON/OFF) expression patterns of two TFs, *bi* and *grn*, control the formation of four LoP layers and corresponding T4/T5 axonal wiring patterns.

## Discussion

Single-cell transcriptional profiling has the potential to transform our understanding of the genetic programs controlling wiring in complex nervous systems (Li et al. 2017; Tasic et al. 2018; Klinger et al. 2018). However, neurons exhibit a vast diversity of wiring patterns, morphologies, and molecular identities, making it difficult to extract the transcriptional logic underlying specific wiring features. Here, we turned to the closely related T4/T5 subtypes of the *Drosophila* visual system, which differ by specific variations in wiring, with the expectation that transcriptional differences among them would reflect the specificity of dendritic inputs and axonal outputs. A unique attribute of T4/T5 neurons is that the same dendritic and axonal wiring patterns are reiteratively used among different subtypes; each neuron can be described by a unique combination of one of four types of axonal outputs and one of two types of dendritic inputs. We anticipated that this property of T4/T5 neurons would provide an opportunity to assess the relationship between specific genetic programs and fundamental features of neuronal architecture.

Unsupervised analysis revealed that separable transcriptional programs correlate with these specific wiring features. We demonstrate through gain and loss of function experiments the functional significance of these programs. These programs can be re-assembled in a modular fashion to generate neuronal subtypes with different combinations of wiring features. A modular transcriptional architecture may provide a general strategy for discrete modifications to neuronal connectivity in development and evolution.

A common T4/T5 neuronal identity is defined by a unique combination of TFs expressed in all subtypes (e.g. *Lim1*, *Drgx*, *acj6*) (Davie et al., 2018; Konstantinides et al., 2018). Perturbation of TFs expressed in all subtypes disrupts overall organization of T4/T5 neurons, including both dendritic and axonal morphologies (Contreras et al., 2018; Schilling et al., 2019). We find that this common T4/T5 transcriptional program is further diversified by separable feature-specific transcriptional programs. These programs are defined by three binary (ON/OFF) TF expression patterns, with two TFs defining the four axonal outputs and one TF defining dendritic inputs.

Four pairs of T4/T5 subtypes with shared axonal outputs (and different dendritic inputs) each target one of four LoP layers, a-d. The ultimate layered architecture of neuropils develops through sequential lamination into increasing numbers of layers (Sanes and Zipursky, 2010; Millard and Pecot, 2018). Together with previous results, our findings suggest that the lamination of T4/T5 axonal outputs occurs via two distinct processes, each controlled by a separate TF. Binary expression of *bi* is required for lamination of the broad a/b from c/d LoP domains (Apitz and Salecker, 2018), whereas binary expression of *grn* is required for sublamination of each domain into separate LoP layers. Importantly, perturbation of each TF exclusively disrupts the corresponding lamination step, while not affecting other morphological features of T4/T5 neurons. Similarly, two quartets of subtypes with shared dendritic inputs (and different axonal outputs) were defined by binary expression of *TfAP-2*. Arborization of dendrites in M10 (T4) or Lo1 (T5) occurs during initial neurite guidance steps, preceding the developmental stages covered in this study (Pinto-Teixeira et al., 2018). We hypothesize that DEGs between T4 and T5 subtypes identified in our analysis contribute to the connections with two distinct sets of presynaptic partners (Shinomiya et al., 2019).

The binary expression patterns of TFs also mirror the developmental lineages of T4/T5 neurons. a/b and c/d subtypes arise from *bi*- and *bi*+ progenitor populations. Neuroblasts from each population undergo two terminal Notch-dependent asymmetric divisions to give rise to the eight subtypes (Pinto-Teixeira et al. 2018). These divisions correspond to binary expression patterns of *grn* and *TfAP-2*, respectively, which act with Notch signaling to regulate wiring. Remarkably, despite divergent developmental trajectories separated by multiple divisions and distinct progenitor pools, all T4 and all T5 subtypes converge onto the same transcriptional programs associated with two types of dendritic inputs. Three regulatory dichotomies could also reflect the evolutionary origin of T4/T5 subtypes and correspond to consecutive duplications of ancestral cell types and circuits (Shinomiya et al. 2015; Arendt, 2016).

Each axonal and dendritic transcriptional program is characterized by a specific pattern of TFs, as well as a set of CSPs, many of which are implicated in regulating wiring in other developmental contexts. These include Ig superfamily proteins in which different paralogs exhibit discrete heterophilic binding specificities, including the beat/side and the dpr/DIP interacting protein families (Zinn and Ozkan, 2017). Interestingly, dynamic expression of these proteins in neurons with shared wiring features was developmentally coordinated. We envision that the synaptic specificity of T4/T5 dendrites and axons are determined by the combined activity of these recognition molecules through interactions with synaptic partners. Future experiments utilizing gain and loss of function analysis, either alone or different combinations, will provide insights into the cellular recognition mechanisms by which synaptic specificity is established.

The composite morphological properties of T4/T5 subtypes allowed us to identify, and thus decouple transcriptional programs for dendrite and axon wiring. Combining separate dendritic and axonal programs, and variations on them, may contribute to the diversification of synaptic specificity in different neuronal subtypes across complex nervous systems

## Methods

### Animal husbandry

Flies (Drosophila melanogaster) were reared at 25°C on standard medium. For developmental analysis by immunohistochemistry, sorting, and sequencing, white pre-pupae (0h APF) were collected and incubated for indicated number of hours.

### Fly Stocks

Multiple-transgene genotypes are enclosed in brackets. The following transgenic lines were used in this study: pBPhsFlp2::PEST (gift from Aljoscha Nern and Gerald Rubin), 10XUAS-IVS-myr::tdTomato (Bloomington Drosophila Stock Center (BDSC #32222)), UAS-FSF-smGdP::HA_V5_FLAG (gift from Aljoscha Nern and Gerald Rubin), 23G12-Gal4 (BDSC #49044), {R59E08-p65ADZp (attP40); R42F06-ZpGdbd (attP2)} (JRC_SS00324, Aljoscha Nern and Gerald Rubin), UAS-H2A::GFP (Barret Pfeiffer and Gerald Rubin), UAS-CD4-tdGFP (BDSC #35839), 23G12-LexA (BDSC #65044), LexAop-myr::tdTomato (Zipursky laboratory), 10XUAS-myr::GFP (Zipursky laboratory), 10XUAS-IVS-mCD8::RFP (BDSC #32219), Mi{PT-GFSTF.1}klg[MI02135-GFSTF.1] (BDSC #59787), Mi{PT-GFSTF.1}beat-IV[MI05715-GFSTF.1] (BDSC #66506), dpr2-Gal4 (Zipursky laboratory), P{w[+mW.hs]=GawB}grn[05930-GAL4] (BDSC #42224), Mi{y[+mDint2]=MIC}beat-VI[MI13252] (BDSC #58680), Mi{PT-GFSTF.0}TfAP-2[MI04611-GFSTF.0] (BDSC #61776), P{y[+t7.7] v[+t1.8]=TRiP.HMS01085}attP2 (UAS-grn-RNAi) BDSC #33746), UAS-grn.ORF.3xHA (FlyORF #F001916), 42F06-Gal4 (BDSC #41253), UAS-p35 (BDSC #5072).

For visualization of T4a clone, virgin females {pBPhsFlp2::PEST; 10XUAS-IVS-myr::tdTomato; UAS-FSF-smGdP::HA_V5_FLAG / CyO::TM6B} were crossed to males with the T4/T5-specific Split-Gal4 driver {R59E08-p65ADZp (attP40); R42F06-ZpGdbd (attP2)} (JRC_SS00324). White pre-pupae were heat shocked at 37°C for 3 minutes. For FACS sorting of GFP+ T4/T5 neurons, 23G12-Gal4 was used to drive UAS-H2A::GFP. For T4/T5 developmental timecourse, 23G12-Gal4 was used to drive UAS-H2A::GFP and 10XUAS-IVS-mCD8:RFP. For visualization of all T4/T5 neurons with subtype-specific markers, female virgins of the genotypes: {23G12-LexA; LexAop-myr::tdTomato; 10XUAS-myr::GFP / CyO::TM6B} or {w; 10xUAS-IVS-mCD8::RFP; 23G12-Gal4} were crossed to males with MiMICs or their derivatives for klg, Beat-IV, dpr-2, Beat-VI, or to w[1] males for Fas2 immunolabeling. For visualization of grn-expressing neurons, grn-Gal4 was used to drive 10XUAS-myr::GFP. For grn phenotypes, virgin females {w[1];UAS-CD4-tdGFP;23G12-Gal4} were crossed to males with UAS-grn-RNAi or UAS-grn.ORF.3xHA transgenes. For p35 rescue experiments, virgin females with 23G12-Gal4, UAS-CD4-tdGFP and with or without UAS-grn.ORF.3xHA were crossed to males with 42F06-Gal4, and with or without UAS-p35 transgene, as indicated. All RNAi and overexpression crosses were raised at 29°C.

### Immunohistochemistry / Immunofluorescence

Brains were dissected in ice-cold Schneider’s Drosophila Medium (Gibco #21720-024), and fixed in PBS (Bioland Scientific LLC #PBS01-03) containing 4% paraformaldehyde (Electron Microscopy Sciences, Cat#15710) for 25 min at room temperature (RT). Brains were rinsed repeatedly with PBST (PBS containing 0.5% Triton-X100 (Sigma #T9284)), and incubated in blocking solution (PBST containing 10% Normal Goat Serum (Sigma #G6767)) for at least 1hr at RT prior to incubation with antibody. Brains were incubated sequentially with primary and secondary antibodies diluted in blocking solution overnight at 4C, with at least 2 PBST rinses followed by 2-hour incubations at RT in between and afterwards. Brains were transferred to 50% (for 30 minutes), then 100% EverBrite mounting medium (Biotium #23001) and mounted on slides for confocal microscopy.

Primary antibodies and dilutions used in this study were as follows: chicken anti-GFP (abcam #13970, 1:1000), rabbit anti-dsRed (Clontech #632496, 1:200), mouse anti-Brp (nc82 from Developmental Studies Hybridoma Bank (DSHB), 1:30), mouse anti-Fasciclin II (1D4 from DSHB, 1:20), mouse anti-V5 (abcam #ab27671, 1:300), rabbit anti-Dcp-1 (Cell Signaling Technology #9578, 1:50). Secondary antibodies and dilutions used in this study were as follows: goat anti-chicken Alexa Fluor 488 (AF488) (Invitrogen #A11039, 1:200), goat anti-mouse AF488 (Invitrogen #A11029, 1:500), goat anti-rabbit AF568 (Invitrogen #A11011, 1:200), goat anti-rat 568 (Invitrogen #A11077, 1:500), goat anti-rabbit AF647 (Invitrogen #A27040, 1:200), and donkey anti-mouse Cy5 (Jackson ImmunoResearch #715-175-150, 1:200).

### Confocal Microscopy and Image Analysis

Immunofluorescence images were acquired using a Zeiss LSM 880 confocal microscope with Zen digital imaging software. Optical sections or maximum intensity projections were level-adjusted, cropped and exported for presentation using Image J software (Fiji). Reported expression patterns were reproducible across three or more biological samples. For cell number quantifications, optic lobes were mounted with ventral side facing objective (as in Figure 1), and a single optical section per lobe was acquired at 3/8 total depth in z-dimension through M10. For quantification of apoptosis, optic lobes were mounted with posterior side facing objective, and a superficial optical section with approximately 300 T4/T5 cell bodies was acquired per lobe. The section depth was determined with Dcp-1 immunofluorescence channel turned off. Files were randomized, and cell numbers and proportion of apoptotic cells were quantified blind to condition using Fiji.

### Single-cell transcriptome profiling

#### Purification of genetically labelled T4/T5 neurons

Males with 23G12-Gal4 driver were crossed to virgin females with UAS-H2A::GFP reporter. F1-generation female white pre-pupae were collected at 0h APF and reared at 25°C. Optic lobes were dissected out at 24h and 48h APF from 27 and 18 pupae, respectively. Brain tissue was incubated in papain (Worthington #LK003178) and Liberase protease (Sigma-Aldrich #5401119001) cocktail at 25°C for 15 min. Next, tissue was gently washed twice with 1X PBS and dissociated mechanically by pipetting. Cell suspension was filtered through 20 μm cell-strainer (Corning #352235). Single-cell suspension was FACS sorted (BD FACSAria II) to isolate GFP-positive cells.

#### Single-cell library preparation and sequencing

FACS-sorted single-cells were captured from a cell suspension using the 10X Chromium platform (~6500-7000 cells loaded). Single-cell RNA-Seq libraries were generated using Chromium Single Cell Reagent Kit V2 according to the manufacturer’s protocol, with 12 cycles of PCR for cDNA amplification. RNA-Seq libraries were sequenced using Illumina Hiseq 4000 platform (paired-end 100 bp reads). Each sample was captured and sequenced using one lane of 10X Chromium and one lane of HiSeq 4000.

### Raw data processing

Raw Illumina base call files (*.bcl files) were converted into fastq files using bcl2fastq (--use-bases-mask=Y26n*,I8n*,Y100n*). Fastq files were processed using Cell Ranger (2.2.0) pipeline with default parameters. Reference transcriptome package for Cell Ranger was generated using *Drosophila melanogaster* genome sequence and gene annotations from FlyBase (release 6.22). Both samples were sequenced at mean depth of 92,000 reads per cell (92% saturation). Average fractions of reads uniquely (confidently) mapped to genome and transcriptome were 93% and 83%, respectively.

### Single-cell data analysis

All steps of single-cell data analysis were performed using functions and methods implemented in Seurat package (2.3.4) (Butler et al., 2018). Analysis for 24h and 48h datasets were performed separately.

#### Quality control and data pre-processing

For 24h dataset, we recovered 3833 cells (median of 1447 genes and 3353 transcripts per cell). For 48h dataset, we recovered 3894 cells (median of 1633 genes and 4389 transcripts per cell). Initial set of cells was pre-filtered based on total number of detected genes (min. 1000; max. 2000), and percentage of mitochondrial transcripts (max. 5%). After pre-filtering, 3312 and 3620 cells remained for 24h and 48h datasets, respectively. Raw transcript counts were log-normalized using NormalizeData function. Next, we regressed out total number of transcripts per cell (nUMI) and scaled expression values to Z-scores using ScaleData function.

#### Preliminary dimensionality reduction and detection of outlier cells

Sets of highly variable genes were selected using FindVariableGenes function (x.low.cutoff: 0.1, x.high.cutoff: 5, y.cutoff: 0.5). Highly variable genes were used to perform independent component analysis (ICA) using RunICA function. Independent components (ICs) were manually inspected to identify and flag outlier cells. In total, 241 and 60 cells were flagged as outliers in 24h and 48h datasets, respectively. Outlier cells were removed from subsequent steps of the analysis. After quality control and filtering, final datasets included 3071 cells for 24h, and 3557 cells for 48h datasets.

#### Dimensionality reduction and clustering (48h APF)

We repeated selection of highly variable genes on final datasets using the same parameters (2290 genes), and used them to perform ICA. Inspection of results of ICA revealed that the three ICs separated cells into two discrete populations of approximate halves. Final clustering was performed based on these 3 ICs using the graph based clustering approach implemented in FindClusters function with default parameters. In addition to ICA, we performed principal component analysis (PCA) using the same set of highly variable genes. Comparison of ICA and PCA results revealed robustness of clusters identified by both methods (Supplementary Figure 1).

t-distributed stochastic neighbor embedding (tSNE) was used to visualize cellular heterogeneity based on ICA and PCA results using RunTSNE function (perplexity: 100). Clusters were validated and matched to eight morphological T4/T5 subtypes using *in vivo* expression patterns of marker genes (Figure 3).

#### Dimensionality reduction and clustering (24h APF)

Similar to 48h dataset, we selected highly variable genes (2198 genes), and used them to perform ICA. We selected three ICs that were driven by similar sets of genes as ICs used for clustering of 48h dataset. Selected ICs were used to perform clustering and tSNE using same parameters as 48h dataset. Cluster identities were matched with T4/T5 subtypes using expression patterns of the same sets of marker genes (Supplementary Figure 4). In comparison to 48h dataset, differences between subtypes at 24h were less pronounced. This may reflect a lower degree of transcriptional divergence among distinct subtypes at earlier stages of development.

#### Differential gene expression analysis

Differentially expressed genes (DEGs) were identified using Wilcoxon rank-sum test implemented in FindMarkers function (min.pct: 0.25, min.diff.pct: 0.25; fold-change: 1.5). We performed this analysis for each cluster against all other cells in dataset (“one versus all”, Figure 1), and between pairs of individual clusters (“one versus one”, Figure 4 and Supplementary Figure 3).

### Data availability

Raw sequencing data (fastq-files), single-cell expression matrix and cell clustering results were deposited to NCBI Gene Expression Omnibus (GEO) under accession: GSE126139.

## Acknowledgments

We thank members of the Zipursky lab, Joshua Sanes, Karthik Shekhar, and Jonathan Flint for critical discussion of the manuscript. We thank Gerald Rubin, Aljosha Nern, Orkun Akin, and Alain Garces for transgenic fly lines and antibodies. We thank Donghui Cheng and Owen Witte for assistance in FACS purification of cells, and UCLA TCGB and BSCRC BioSequencing Core Facility for assistance with single-cell RNA-Sequencing. This work was supported by the NIH National Institute of Neurological Disorders and Stroke (T32NS048004) (S.A.L), and the G. Harold and Leila Y. Mathers Foundation (S.L.Z.). S.L.Z. is an Investigator of the Howard Hughes Medical Institute.

## Author contributions

Y.Z.K., J.Y., S.A.L., and S.L.Z. conceived the project and designed the experiments. Y.Z.K and J.Y. generated single-cell sequencing data. Y.Z.K performed single-cell data analysis. J.Y. and S.A.L performed expression validation experiments. S.A.L. performed functional experiments. Y.Z.K., J.Y., S.A.L., and S.L.Z. interpreted results and wrote the manuscript.

---

**Supplementary Figure 1.**
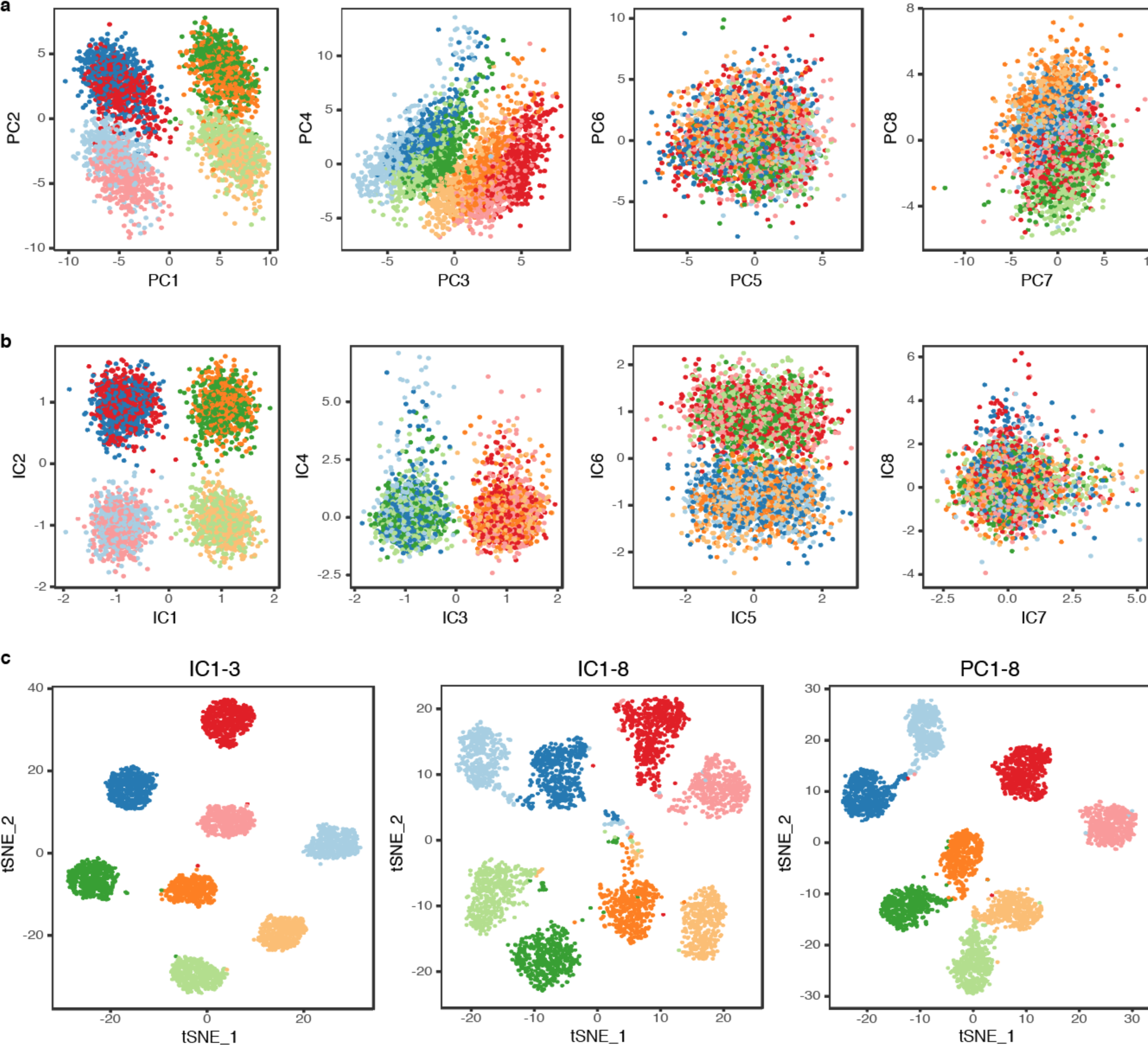
T4/T5 neurons robustly cluster into eight transcriptionally distinct populations (48h APF). (a) Principal component analysis (PCA). (b) Independent component analysis (ICA). Distributions of cells along eight principal components (PCs) and eight independent components (ICs) (b). (c) tSNE plots on IC 1-3 (left), IC 1-8 (middle), PC 1-8 (right). Cells are color coded according to the final clustering results based on IC 1-3, as in Figure 1.

**Supplementary Figure 2.**
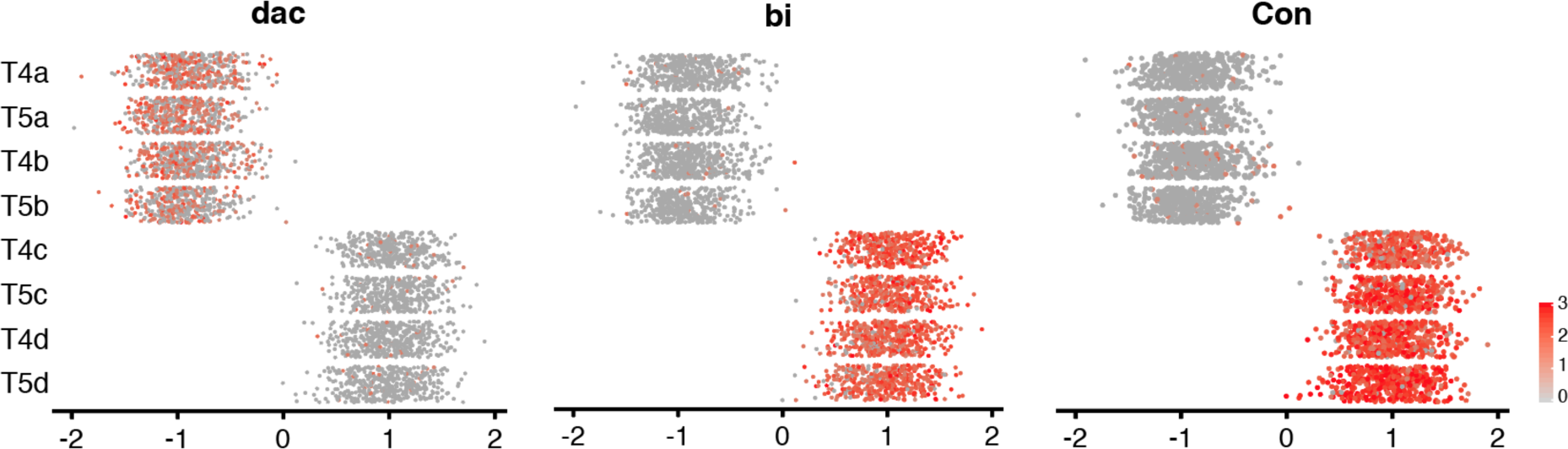
Expression patterns of known marker genes for a/b and c/d subtypes along Axis 1 at 48h APF. See legend of Figure 3 for details.

**Supplementary Figure 3.**
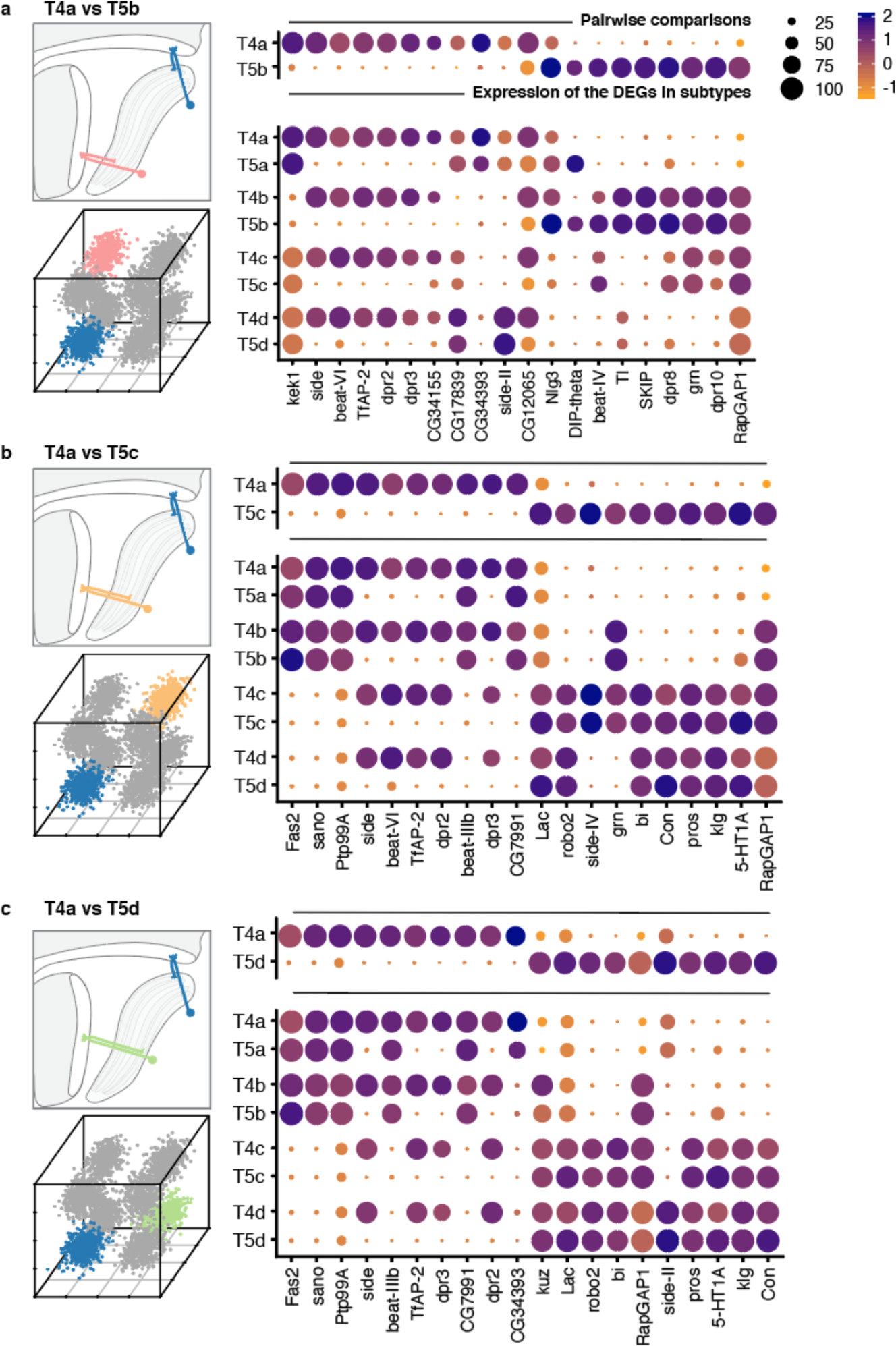
Pairwise comparisons between T4a and subtypes that differ by both axonal outputs or dendritic inputs. Top 20 DEGs are shown for each comparison. See legend of Figure 4 for details.

**Supplementary Figure 4.**
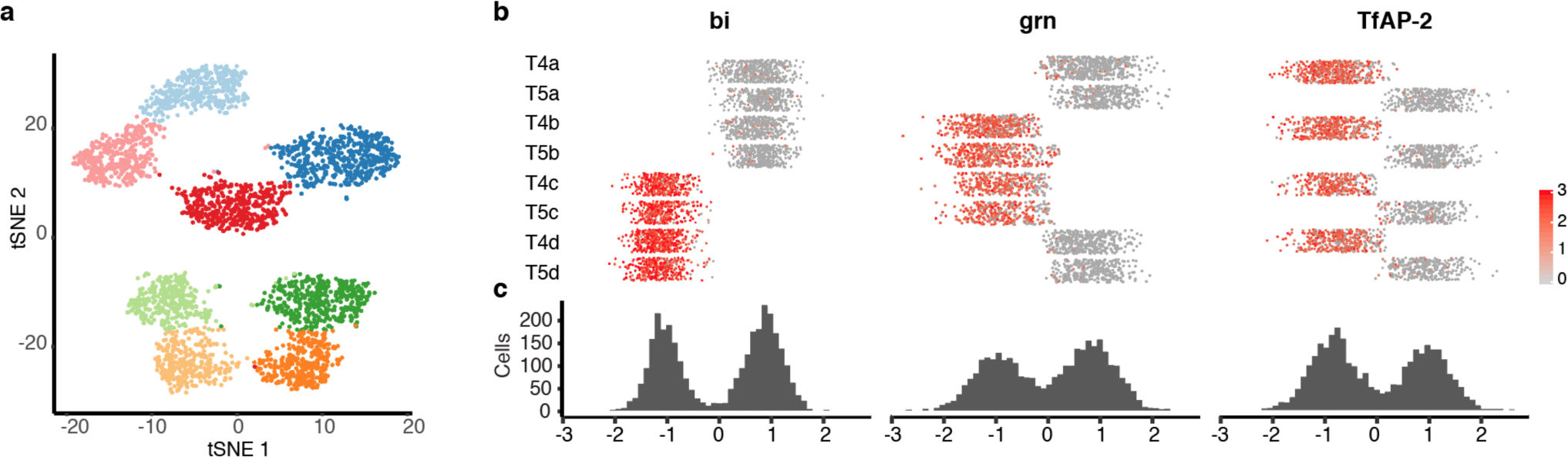
Single-cell profiling of T4/T5 neurons at 24h APF. Unsupervised analysis revealed eight transcriptionally distinct populations. (a) tSNE plot of 3833 single-cell transcriptomes. (b). Distribution of cells along three primary axes of transcriptional diversity, and expression patterns of TF with highest contribution to each axis. See legends of Figures 1-3 for details.

**Supplementary Figure 5.**
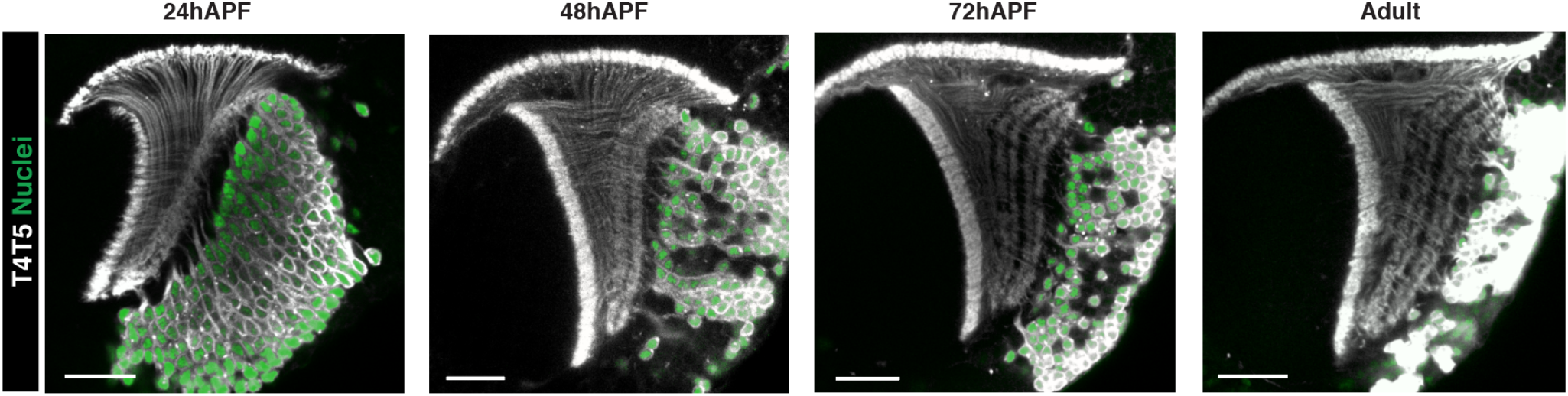
Sequential lamination of T4/T5 axons and four LoP layers. 23G12-Gal4 drives membrane localized RFP (grey) and nuclear localized GFP (green) in all T4/T5 neurons throughout pupal development.

**Supplementary Figure 6.**
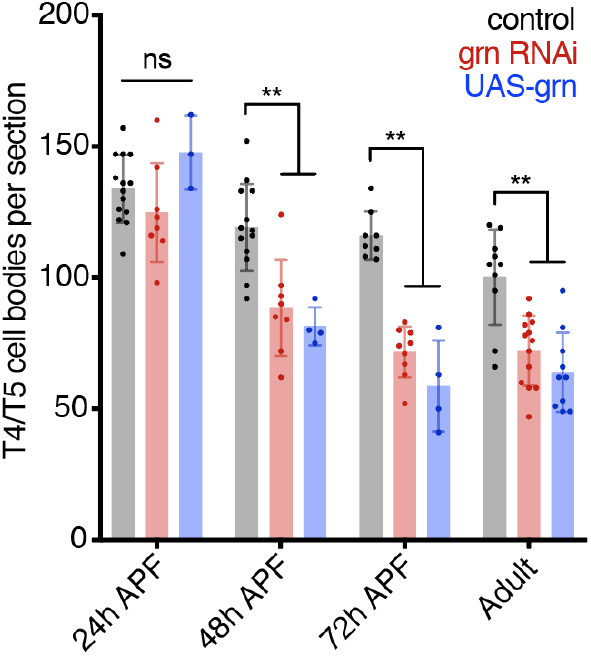
*grn* RNAi and *grn* overexpression cause significant loss of T4/T5 neurons between 24 and 48h APF. Statistical significance assessed by one-way ANOVA with Dunnett’s multiple comparison test (**p<0.01).

